# GiraFFe Browse: A lightweight web based tool for inspecting GFF and FASTA data

**DOI:** 10.1101/273631

**Authors:** Owen Garland, Amanda Clare, Wayne Aubrey

## Abstract

**Summary:** GiraFFe Browse is a tool aimed at making GFF and FASTA data more useful and accessible to researchers. Existing solutions are large in scope and difficult to maintain without specialist knowledge of computer systems. GiraFFe Browse is a lightweight alternative, built using modern software engineering practices with a focus on being developer and user friendly.

**Availability and implementation:** GiraFFe Browse is open source (GPL) software that is available from the Github repository: https://github.com/bag-man/giraffe-browse

**Demo version:** An example installation of the application is available at http://giraffe.owen.cymru, using a GFF and FASTA file of *Escherichia coli* from the European Nucleotide Archive.

## Introduction

DNA sequencing is now routine for a growing number of research laboratories due to the current high throughput sequencing instruments available [3]. Making sense of this data has been made easier with the introduction of new genome annotation pipelines such as Prokka [7], which identify and label features of a DNA sequence, and produce output in a GFF3 format [2]. However, the problem of examining these annotations and making them accessible is still challenging. Researchers lack a user friendly interface for querying and interacting with their data, that is simple to setup without much specialist knowledge of systems administration.

Existing platforms for storing and interacting with genomic data include GMOD, InterMine, Ensembl and IGV [9, 4, 11, 10]. The Generic Model Organism Database Project (GMOD) is a large consortium of tools with a very wide set of features. At its core is Chado [5], a large PostgreSQL database schema, aimed at being generic enough for all possible genomes to be represented within it. While this solution is suitable for large projects, due to the nature of trying to be a “one size fits all” solution, it often comes with too many features and options for projects with smaller scope. This can make it unsuitable for projects with short term analysis and a lack of expertise in systems administration, as the time taken to setup GMOD may be longer than the time needed to analyse the data. The GMOD project provides JBrowse [8], a Javascript/HTML5 genome browser, which requires BioPerl and other Perl modules. This provides a scrolling view of a genome useful for those who want distribution overviews, or to inspect the region surrounding a gene of interest. GMOD also provides Tripal [6], a web interface to a Chado database based on the Drupal CMS, allowing editing. InterMine is a data warehouse for the integration and querying of biological data. It can integrate Chado databases, GFF and FASTA files and more sources, and now powers the online databases/websites for many of the major model organisms. Ensembl is another option similar to GMOD in its scale, based on Perl and SQL, but for vertebrate genomes. GenomeHubs [1] aims to make the installation and setup of Ensembl easier by offering a containerised solution. This is a large improvement on the accessibility of the software, however it is still aimed at the larger long term projects. Interactive Genome Viewer (IGV) has far fewer installation requirements. It is a Java application for viewing the contents of BAM/BED/GFF/many other file formats. It provides a zoomable, scrollable track-level view, from whole genome down to individual bases, and will show alignments along with genome annotation features.

Even though these tools offer many features, visualisations and analysis of data, their barriers to entry can result in sequences and annotations persisting in FASTA and GFF files, not being fully exploited. We have created a tool that allows researchers to have an easy and flexible way to extract meaningful information from their annotation pipelines, as well as making the data more accessible to colleagues. The application has two core parts: a script that will ingest the FASTA and GFF data files into a MongoDB NoSQL database, and a web interface that allows researchers to browse and search that database for information relevant to their research.

## Features

### Simple installation and maintenance

GiraFFe Browse is built with developer and user friendliness in mind. GiraFFe Browse is built with Node.js and uses the Node Package Manager to install and manage dependencies. This means that the installation and maintenance of all the required libraries for a project can be done with one simple command. All dependencies are installed local to the application folder, and not system wide. This allows multiple applications to be installed alongside one another using differing versions of the same packages without version and dependency conflicts. Uninstalling is a simple case of deleting the application directory, rather than uninstalling system wide dependencies.

To streamline the installation process an automated script has been provided for OS X and Debian/Ubuntu systems that will install the required dependencies. Once installed the user can either manually run the web server on a port of their choosing, or on systemd based systems, install the project to run as a service on the host machine, ideal for shared servers with multiple users. With the choice of Node.js, and the easy installation script, GiraFFe Browse is well suited to short term projects and researchers who need to quickly inspect genome annotations.

### NoSQL database solution with MongoDB

After installation, the researcher will import the sequence and annotation data, provided in the form of a FASTA and GFF3 files. Those datasets are parsed into JSON format, and stored in a NoSQL database in two collections. One is simply a JSON representation of the FASTA file. The other stores a document for each record in the GFF3 file, as well as adding extra fields, including the corresponding nucleotide sequence extracted from the coordinates provided in the GFF3 file.

The database is independent of the web application and can be be accessed through the MongoDB shell for more complex queries by advanced users if required. This also has the advantage of allowing scope for developers to create tools based on this data structure.

### Automatic retrieval of coding sequences and protein sequences

GFF3 is a common format for sequence annotation that includes the coordinates of the start and the end of an annotated sequence, along with the details of the annotation. GiraFFe Browse extracts the sequence from the corresponding contig, in the correct reading frame. It also provides reverse complement and protein coding translations, with a handy copy-to-clipboard button.

### Filterable and queryable

The GFF3 file format contains nine fields. The ninth field (‘attributes’) is a flexible list of key-value pairs, designed to hold user-defined annotations. While this is very flexible for researchers who need to add their own metadata, applications that present this data need to be equally flexible.

The GiraFFe Browse web interface displays a list of records in the GFF3 file, showing the contents of the fields present in all GFF3 files. Due to the flexibility of the attributes field, the user can also choose the extra fields they wish to see from a series of pre-populated drop down lists of the fields that exist in the GFF file. Once these fields have been selected, a search can be performed for records that have attributes matching a search query. If multiple fields are selected, the query uses each field as an AND selector, which means that only results that contain data in all of the selected columns will be returned.

There is also an option to filter results by the type of sequence that has been annotated in the GFF3 file, for example mRNA, gene, or CDS. These types are constrained to be terms from the Sequence Ontology [2].

### Use cases

- A lecturer would like to make the *E. coli* genome annotations available to their class in a browseable online format in order that each student can each query for transposases, copy the sequences and perform subsequent analyses.
- A research group would like to confirm that a newly sequenced strain of *E. coli* is functionally similar to a previously sequenced wild-type and wish to quickly inspect the genome for the presence or absence of certain genes.
- An environment is routinely sampled for metagenomic sequencing to test for the presence of antibiotic resistant genes. The annotated sequences need to be quickly viewed to support a timely decision.

## Discussion

The advent of cheap sequencing technology is democratising the process of collecting and interpreting genomes and sequencing will soon become a transient and disposable commodity. Genome science is no longer restricted to large institutions but is now available for small laboratories and groups who specialise in the study of certain organisms. After assembly and annotation of the sequence data, these researchers need to be able to explore the genomes using tools that are easy to use, easy to install and allow group read access.

GiraFFe Browse has minimal dependencies. The core only requires MongoDB to store the data and Node.js to run the webserver that provides the web interface. These widely used software packages are well supported on many platforms.

Future developments will include offering an externally hosted version for those who prefer no installation, and further search, sorting and filtering options.

## References

[1] R. J. Challis et al. GenomeHubs: simple containerized setup of a custom Ensembl database and web server for any species. Database, 2017(1):bax039, 2017.

[2] K. Eilbeck et al. The Sequence Ontology: A tool for the unification of genome annotations. Genome Biology, page 6:R44, 2005.

[3] S Goodwin, J. D. McPherson, and W. R McCombie. Coming of age: ten years of next-generation sequencing technologies. Nature Reviews Genetics, 17(6):333–351, 2016.

[4] A. Kalderimis et al. InterMine: extensive web services for modern biology. Nucleic Acids Research, 42:W468–72, 2014.

[5] C. J. Mungall et al. A Chado case study: an ontology-based modular schema for representing genome-associated biological information. Bioinformatics, 23(13):i337–i346, 2007.

[6] L-A. Sanderson et al. Tripal v1.1: a standards-based toolkit for construction of online genetic and genomic databases. Database, 2013, 2013.

[7] T. Seemann. Prokka: rapid prokaryotic genome annotation. Bioinformatics, 30(14):2068–2069, 2014.

[8] M. E. Skinner, A. V. Uzilov, L. D. Stein, C.J. Mungall, and I.H. Holmes. JBrowse: A next-generation genome browser. Genome Research, 19(9):1630–8, 2009.

[9] L. D. Stein et al. The generic genome browser: a building block for a model organism system database. Genome Research, 12(10):1599–610, 2002.

[10] H. Thorvaldsdéottir et al. Integrative genomics viewer (IGV): high-performance genomics data visualization and exploration. Briefings in Bioinformatics, 14:178–192, 2013.

[11] D. R. Zerbino et al. Ensembl 2018. Nucleic Acids Research, 46(D1):D754–D761, 2018.

